# Impact of modular mitochondrial epistatic interactions on the evolution of human subpopulations

**DOI:** 10.1101/505818

**Authors:** Pramod Shinde, Harry J. Whitwell, Rahul Kumar Verma, Mikhail Ivanchenko, Alexey Zaikin, Sarika Jalan

## Abstract

Investigation of human mitochondrial (mt) genome variation has been shown to provide insights to the human history and natural selection. By analyzing 24,167 human mt-genome samples, collected for five continents, we have developed a co-mutation network model to investigate characteristic human evolutionary patterns. The analysis highlighted richer co-mutating regions of the mt-genome, suggesting the presence of epistasis. Specifically, a large portion of COX genes was found to co-mutate in Asian and American populations, whereas, in African, European, and Oceanic populations, there was greater co-mutation bias in hypervariable regions. Interestingly, this study demonstrated hierarchical modularity as a crucial agent for these co-mutation networks. More profoundly, our ancestry-based co-mutation module analyses showed that mutations cluster preferentially in known mitochondrial haplogroups. Contemporary human mt-genome nucleotides most closely resembled the ancestral state, and very few of them were found to be ancestral-variants. Overall, these results demonstrated that subpopulation-based biases may favor mitochondrial gene specific epistasis.

## 1. Introduction

Genetic polymorphism varies among a species as well as within genomes and carries important implications for the evolution and conservation of species. Polymorphism in the mitochondrial (mt) genome is routinely used to trace ancient human migration routes and to obtain absolute dates for genetic prehistory (Chen, at al., 1995). The human mt-genome is very small (16.6 kb), maternally inherited, evolves in both neutral and adaptive fashions, and shows a great deal of variation as a result of divergent evolution. An absence of recombination within mt-genome provides distinct polymorphic loci which have been used to define human genealogy referred to as mt-genome haplogroups (Chen, at al., 1995). These haplogroups are formed as a result of the sequential accumulation of mutations through maternal lineages. Since mitochondria are essential to cellular metabolism, mt-genome variation has been associated with multiple complex diseases including Alzheimer’s disease in haplogroup U (Van, et al., 2004), idiopathic Parkinson disease within JT haplogroup (Hudson, et al., 2013) and age-related macular degeneration in the JTU haplogroup cluster (Kenney, et al., 2013). Due to population migration, distinct lineages of mt-genome are associated with major global groups (African, American, European, Asian and Oceanic) raising the possibility that mt-genome variation could contribute to the differences in disease prevalence observed among both ethnic and racial groups (Mishmar, et al., 2003; Shriner, and Keita., 2016; Zanellati, et al., 2015).

Conventionally, analyses of mt-genome evolution have focused on individual mutations, particularly in describing haplogroups, and to understand and predict ancestral behavior. However, the evolutionary behavior of mt-genome often involves cooperative changes within and between genes which are difficult to detect using haplogroup analysis. For example, correlated mt-genome mutations were reported among different oxidative phosphorylation subunits, which were found to affect population specific human longevity (Raule, et al., 2014; Fan, et al., 2016; Giuliani, et al., 2018; Conte, et al., 2018). Besides, cooperative activities of both mitochondrial proteins and tRNA genes are critical for mt-genome evolution. The importance of co-mutational interactions has been well documented in the genomics field (Lane, et al., 2012; Chen, et al., 2013; Haddad, et al., 2018). Increasing evidence suggests that interactions among polymorphic sites may confer a cumulative association of multiple mutations with many diseases (Chen, et al., 2013). Interactions among polymorphic sites have also effectively been used to infer ancestry and functional convergence in the human populations (Ioannidis, et al., 2001). Commonly used methods include tree ensembles, functional nodal mutations, and single nucleotide polymorphism (SNP) based enrichment (Lunetta, at al., 2004). Important information about mt-genome evolutionary behavior, which is contained in the correlated changes between nucleotide positions both within and between genes, is not captured by these techniques. Despite strong evidence that mt-genome variation plays a role in the development and progression of complex human diseases, mitochondrial genetic variation has been largely ignored in the context of co-mutations and particularly the mechanisms by which these co-mutations occur (Boles, et al., 1998; Goodman, et al., 2006). Investigation of co-mutation effects can, therefore, improve the explanatory ability of genetics twofold. Firstly, the interaction between two informative genomic positions to explain a part of the trait heritability. Secondly, finding significant statistical links between mutations could provide strong indications of molecular-level interactions that differ between distant populations (Hartwig,, 2013).

Complex network science revolves around the hypothesis that the behavior of complex systems can be elucidated in terms of structural and functional relationships between their constituents employing a graph representation (Albert, and Barabási., 2002; Shinde, et al., 2015; Shinde, and Jalan., 2015; Whitwell, et al., 2018; Rai, et al., 2018; Ho, et al., 2014). The basis of the current study is that genome positions can impact each other and co-mutate within genomes (Shinde, et al., 2018; Du, et al., 2008; Sun, et al., 2014). The interaction between two or more genetic loci is referred to here as the co-mutation of nucleotide positions. There are previous studies which have used genomic co-mutations as a basis of the evolution of human H3N2 and Ebola viruses (Du, et al., 2008; Deng, et al., 2015). These viral genome models have identified the co-mutating nucleotide clusters, apparently underpinning the dynamics of virus evolution since these clusters were antigenic regions of the viral capsid proteins (Du, et al., 2008; Deng, et al., 2015). In another study, Shinde et al. (Shinde, et al., 2018) demonstrated the impact of codon position bias while forming co-mutations using human mt-genomes. These studies have considered perfect co-mutation as causing factor for comutations. However, the role of the co-mutation frequency in these studies remains unclear. Here, we thoroughly examined a set of networks associated with a range of co-mutation frequencies and chose a particular co-mutation frequency for further network construction. Whilst pair-wise co-mutations can be straightforwardly perceived, the identification of larger sized functional units is not straightforward. Here, we used community detection algorithms to enumerate lists of modules formed within networks and described the functional relationships among nucleotide positions forming these modules.

We set out to develop a comprehensive approach to understand mitochondrial diversity using mitochondrial co-mutations. To this end, we conducted a comparative analysis of 24,167 sequenced mt-genomes. The paper is organized as follows. In the first section, we briefly described the level of diversity observed among underlying subpopulations concerning polymorphic site variations in human mitochondrial genomes. In the second section, we described the framework to investigate co-mutations, which are critical in underlying complex mitochondrial evolution. For this, we constructed co-mutation networks which were used to identify modules of co-mutations and also compared these results with those of the corresponding random networks. In the third and fourth sections, we identified local topological phenomena, which were crucial agents for co-mutation networks make-up. We listed down modules comprised of co-mutations and demonstrated that the identified modules indeed correspond to ancestry based associations. Overall, revealing the importance of co-mutational biases among different human subpopulations, our analysis identified local preferences, which were key agents in forming mt-genome epistatic interactions.

## 2. Methods and Material

### 2.1. Acquisition of genomic data

Analysis of mt-genome variations in continental populations has revealed the most ancient of all human continent-specific haplogroups in Africa and their subsequent migration and settlement in other continents (Chen, at al., 1995). Therefore, continents are constituents to defining different global-ancestral lineages beyond being just landmasses. The global lineages among each continent have been shown to explain a variety of signatures including demographic history, climate and environmental changes, local-admixture patterns (Conte, et al., 2018; Fonseca, et al., 2008; Derenko, et al., 2001; Hudson, et al., 2013). Each continent has its own signatures as well as shared signatures, as human migrations are known to happen differently among different continents (Mishmar, et al., 2003).

Having this notion, we prepared an extensive collection of mitochondrial genomes of geographically diverse *Homo sapiens* populations (Fig 1) from the Human Mitochondrial Database (Hmtdb) (Rubino, et al., 2012). All downloaded genome sequences were in FASTA format. In total, the dataset comprised of 24,167 mitochondrial genome sequences from the five world continents (genome groups), including 3426 African (AF), 2650 American (AM), 8483 Asian (AS), 8060 European (EU) and 1548 Oceanic (OC) genomes. Antarctica was excluded from the present analysis since no data was available. It should be noted that these genome groups are multiethnic cohorts representing a range of populations across the continent (Fig 1A). A brief description of all the genomes and their origin is provided in S1 File.

**Figure 1:**
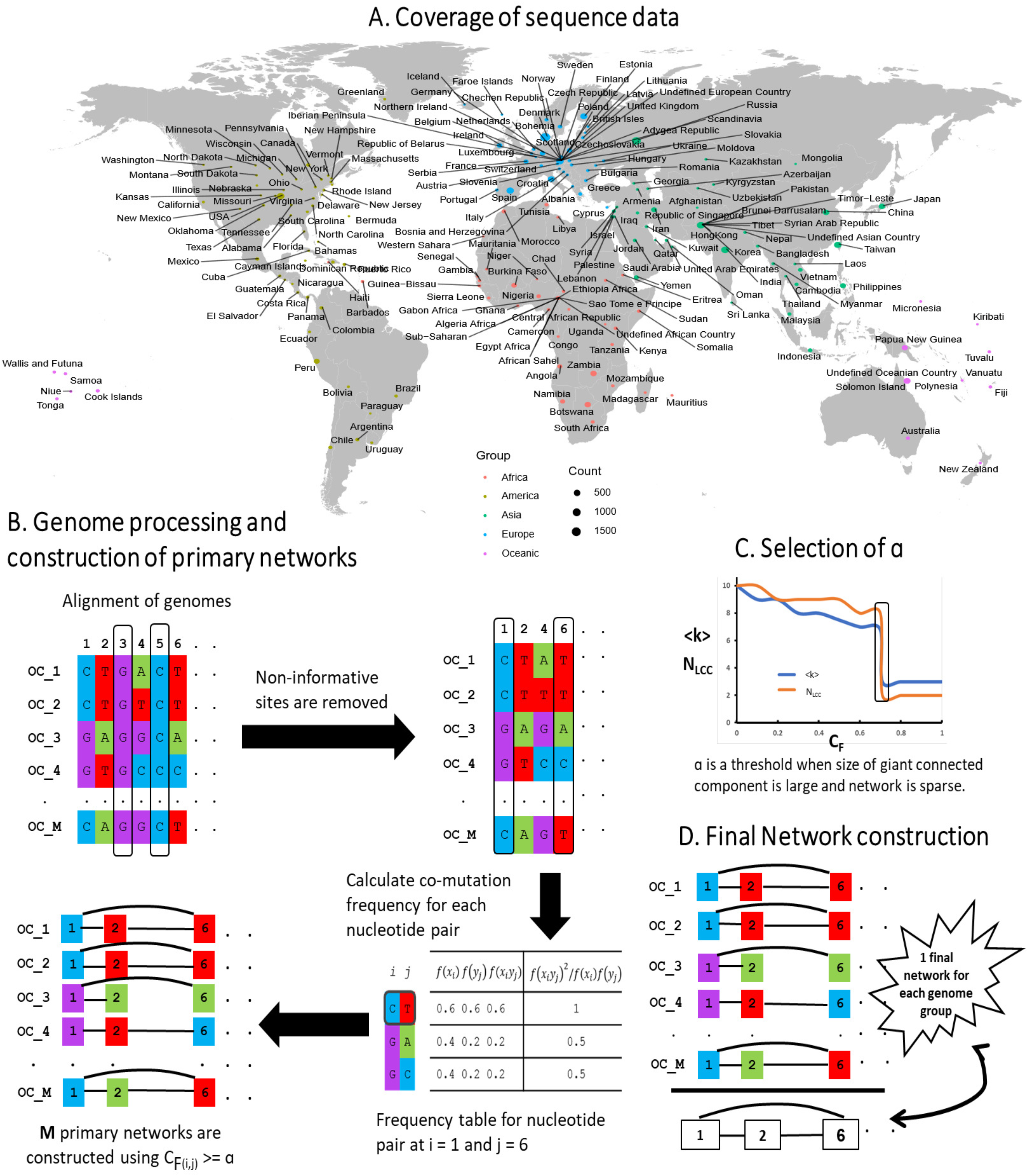
Schematic representation of mtDNA co-mutation network construction and analysis. **(A)** World map shows sequence data taken for the current study covered a good distribution across the entire globe. **(B)** A schematic diagram is drawn for a genome group with 5 sample sequences. The schematic diagram depicts (1) Alignment of genomes. All mitochondrial genomes in a genome group were end to end aligned, and therefore all aligned sequences had the same length. (2) Removal of non-informative sites. A genome position consist of a single nucleotide among all samples was removed from the analysis. (3) Calculation of co-mutation frequency (*C*_*F*_) for each nucleotide pair. **(C)** Selection of network efficiency score (*α*). *α* was a threshold when the average degree (*〈k〉*) of a network is small, and the size of the largest connected component (*N*_*LCC*_) is high. For each genome group, *α* was found to be different. **(D)** Each genome group has *M* genomes *i.e. M* networks. A unique list of edges was_7_picked up from *M* networks from a genome group to construct a final weighted network for *M* networks in a genome group. Likewise, five networks were constructed for five genome groups.

### 2.2. Construction and preliminary analysis of co-mutation networks

Co-mutation calculations were carried out on each genome group distinctly. Co-mutation network construction is broadly divided into two parts, construction of primary networks for each genome sequence followed by the construction of final networks for each genome groups. Each primary comutation network represents an individual sequence, and thus for each genome group, *M* primary co-mutation networks were generated where *M* is the number of sequences in the genome group. In a co-mutation network (for both primary and final), nodes represent genome positions, and edges between nodes represent genomic co-mutations. We constructed five co-mutation networks for each genome group using their primary co-mutation networks. Final co-mutation networks comprise of qualitative information of interactions between genome positions. The methodology for constructing primary and final co-mutation networks is schematically represented in Fig 1 and described as follows:

#### 2.2.1. Primary co-mutation network

(1) Genome sequence data was end to end aligned. (2) All non-variable genome positions within samples of a genome group were removed, leaving only polymorphic genome positions. The number of polymorphic sites (*N*_*P*_) is given in Table 2. (3) Using only polymorphic nucleotide positions, we calculated the frequency of occurrence of all the nucleotide pairs *f* (*x*_*i*_*y*_*j*_) = *N* (*x*_*i*_*y*_*j*_)*/M* where, *N* (*x*_*i*_*y*_*j*_) denoted the number of co-mutation pairs (*x*_*i*_*y*_*j*_) at position (*i, j*). We then calculated the frequency of occurrence of single nucleotides *f* (*x*_*i*_) = *N* (*x*_*i*_)*/M* and *f* (*y*_*j*_) = *N* (*y*_*j*_)*/M* where, *N* (*x*_*i*_) and *N* (*y*_*j*_) denoted the number of single nucleotides at their respective positions *i* and *j* (Du, et al., 2008). (4) Co-mutation of two nucleotides (*C*_*F*_) at position (*i, j*) was denoted as,

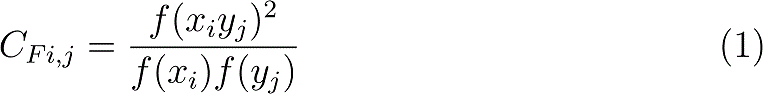

For a particular co-mutation frequency threshold, here, termed as network efficiency score (see Section 2.3), we constructed primary co-mutation networks. A network can be represented mathematically by an adjacency matrix (*A*) with binary entries.

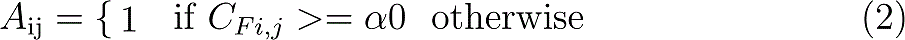

As each genome sequence has its own information of co-mutating genome positions, a total *M* primary co-mutation networks were generated for each genome group.

#### 2.2.2. Final co-mutation network

Unique edges from all primary co-mutation networks of a genome group were used to construct a final co-mutation network (Fig 1D). These five final co-mutation networks were used for network analysis and community detection.

We extracted hierarchical modules from final co-mutation networks and compared these networks with random networks, hypothesizing that hierarchical modularity is the underlying phenomena of co-mutation networks and is not a mere outcome of the random evolutionary process. Furthermore, we characterized the identified module structures using ancestral markers.

#### 2.2.3. Preliminary analysis of co-mutation networks

The degree of a node (*k*_*i*_) is defined as a number of edges connected to the node such as 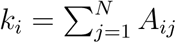 where *N* denoted the number of nodes in a network. The average degree connectivity *(k)* is the average nearest neighbor degree of nodes with degree *k*. The clustering coefficient (C) is a measure of the extent to which nodes in a network tend to cluster together. An average clustering coefficient of a network can be written as 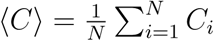. Another property of the network which turns out to be crucial in distinguishing the individual networks was the assortative coefficient (*r*), which measures the tendency of nodes with the similar numbers of edges to connect. The assortative coefficient, *r*, was defined as the Pearson correlation coefficient of degree between pairs of linked nodes (Newman, et al., 2003). The value of *r* being zero corresponds to a random network, whereas the negative (positive) values correspond to dis (assortative) networks.

### 2.3. Selection of network efficiency score (α)

Network efficiency score (*α*) was used to filter edges required for network construction. Selecting an *α* value for each network should require iterating through a range of *C*_*F*_ values. To consider a network with *C*_*F*_ values of least 10^*−*4^ precision would require the construction of 24,167 *∗* 10^4^ networks in total, which would be a very computationally intensive process. Therefore, we performed statistical sampling on each genome group interdependently by selection analysis of *m* samples from each population. The sample size was determined by Cochran’s sample size formula (Cochran,, 1997) with critical value (*z* = 1.96). As the population was finite, the sample size was corrected by Cochran’s adjustment (Cochran,, 1997).

A zero *α* value would result in co-mutation between each mutation and all others, whereas *α* equal to one would give only those pairs of mutations which have co-mutated perfectly in a genome group. In other words, zero *α* value would result in the globally connected network (Fig 2B) and *α* = 1 would result networks with many globally connected small sub-graphs (Fig 2E). Even when the *α* value was as high as 0.99, networks remained very densely connected (Fig 2C). Therefore, it was reasonable to propose a criterion to select an *α* value, otherwise generated networks would be saturated structures holding no information about co-mutations. In order to tackle this, we plotted *〈k〉* and the size of the largest connected component (*N*_*LCC*_) against all the *α* values. We observed surprising network phenomena where at a particular *α* value, *〈k〉* is small whilst *N*_*LCC*_ is large. At this point, networks are sparser as compared to previous *α* values (Fig 2D). By a sparse network, we would mean that the majority of elements of the adjacency matrix are zeroes. After exceeding this *α* value, the network breaks into several disconnected components.

**Figure 2:**
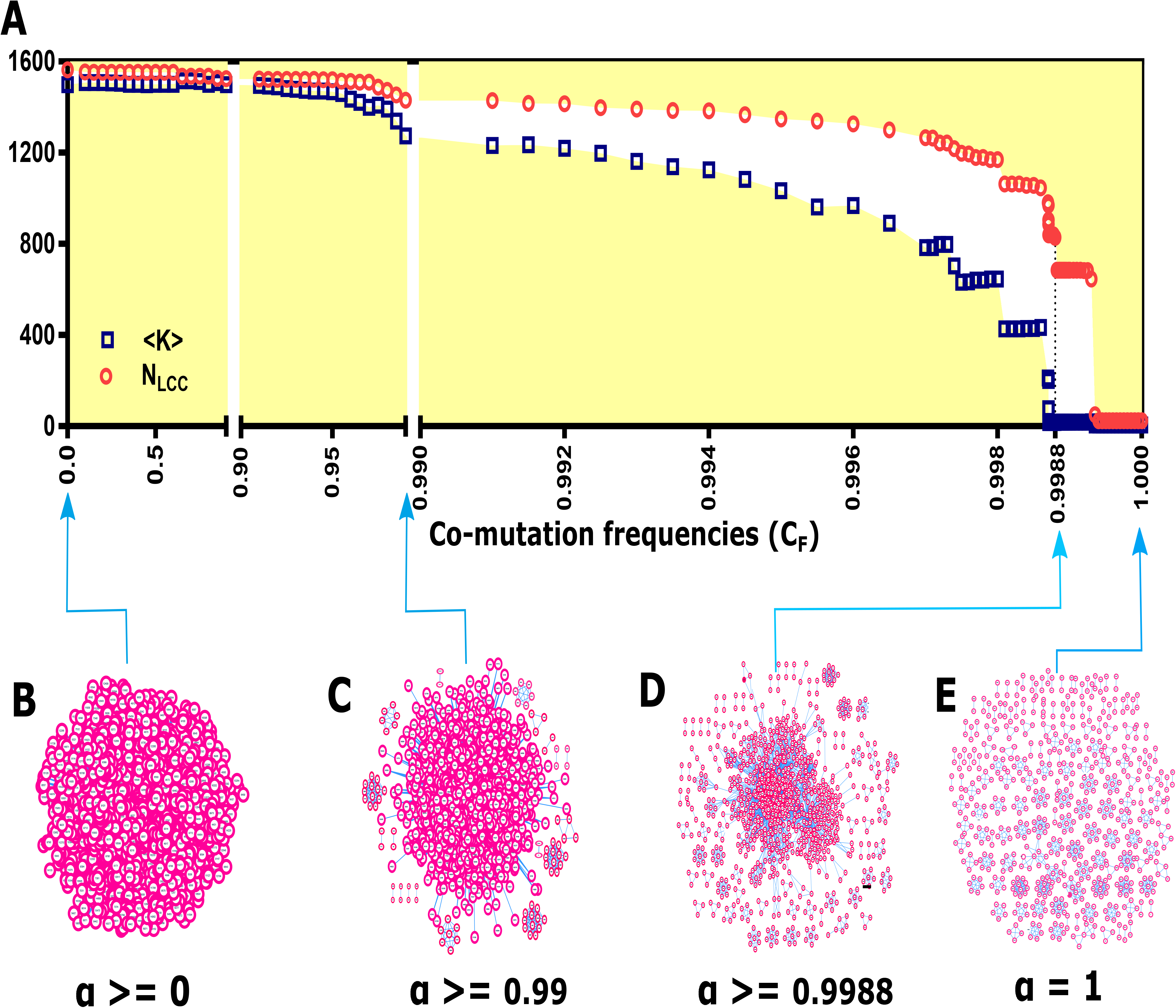
Evolution of mitochondrial co-mutation network. **(A)** The relative size of the largest component and the average degree of the largest component are plotted against co-mutation frequencies (*C*_*F*_). The figure illustrates that at a particular *α* value (for OC, *α* = 0.9988) co-mutation network has both a smaller value of the average degree and the number of nodes in the largest component are sufficiently in large number. We picked this *C*_*F*_ value for network construction. **(B-E)** A sample network at different *C*_*F*_ values show how it evolves from a globally connected network to the network with many disconnected components.

With this criterion, we chose a particular *α* value for each genome group and constructed primary co-mutation networks. Although the *α* value applied to each genome group was very high (close to 1), this value was sufficient to capture more than 50% of the polymorphic sites in each genome group (except in AS; S4 Table and S3 Fig). A similar criterion of filtering network edges has been earlier used to construct gene co-expression network (Jackson, et al., 2018). It should be noted that *α* values for each genome groups were different (Table 2).

### 2.4. Detection of module structures in co-mutation networks

We used the Louvain algorithm, a modularity maximization algorithm, for community detection for our networks (Blondel, et al., 2008). The Louvain method was a simple, efficient, and easy-to-implement method for identifying communities in large networks. The python package of Louvain algorithm was used to enumerate module structures (Blondel, et al., 2008), and Gephi software was used for visualization (Bastian, et al., 2009).

## 3. Results

Analysis of polymorphic sites is provided in supplementary materials and summarised in Table 1 and Fig 3. Codon position (CP) 2 showed fewer polymorphisms as compared to CP 1 and CP 3. Genes *ATP6* and *ATP8* demonstrated a higher level of polymorphisms at all three CPs. Similarly, all three HVS regions have displayed a higher level of polymorphisms, whereas genes of rRNA and tRNA have shown lower levels of polymorphisms.

**Table 1:** Gene- and codon-wise polymorphisms among 13 protein-coding genes. The observed polymorphisms in each of 13 protein-coding genes show mutational biases at codon positions. ATP genes contained the most polymorphisms. CP 2 showed fewer polymorphisms as compare to CP 1 and CP 3. *COX1* and *ND4* had the lowest proportion of observed polymorphic sites, and *ATP6* had the largest proportion.

| Gene Names (OMIM Ids) | Gene (%) | CP 1(%) | CP 2(%) | CP 3(%) |
| --- | --- | --- | --- | --- |
| *ATP6* (516060) | 69 | 69 | 51 | 88 |
| *ATP8* (516070) | 66 | 67 | 54 | 78 |
| *COX1* (516030) | 41 | 25 | 11 | 88 |
| *COX2* (516040) | 47 | 34 | 20 | 87 |
| *COX3* (516050) | 50 | 39 | 26 | 86 |
| *CYB* (516020) | 55 | 49 | 26 | 90 |
| *ND1* (516000) | 47 | 34 | 21 | 87 |
| *ND2* (516001) | 46 | 36 | 19 | 84 |
| *ND3* (516002) | 41 | 33 | 18 | 73 |
| *ND4* (516003) | 42 | 27 | 13 | 87 |
| *ND4L* (516004) | 39 | 27 | 8 | 82 |
| *ND5* (516005) | 47 | 36 | 17 | 88 |
| *ND6* (516006) | 47 | 32 | 21 | 88 |

**Figure 3:**
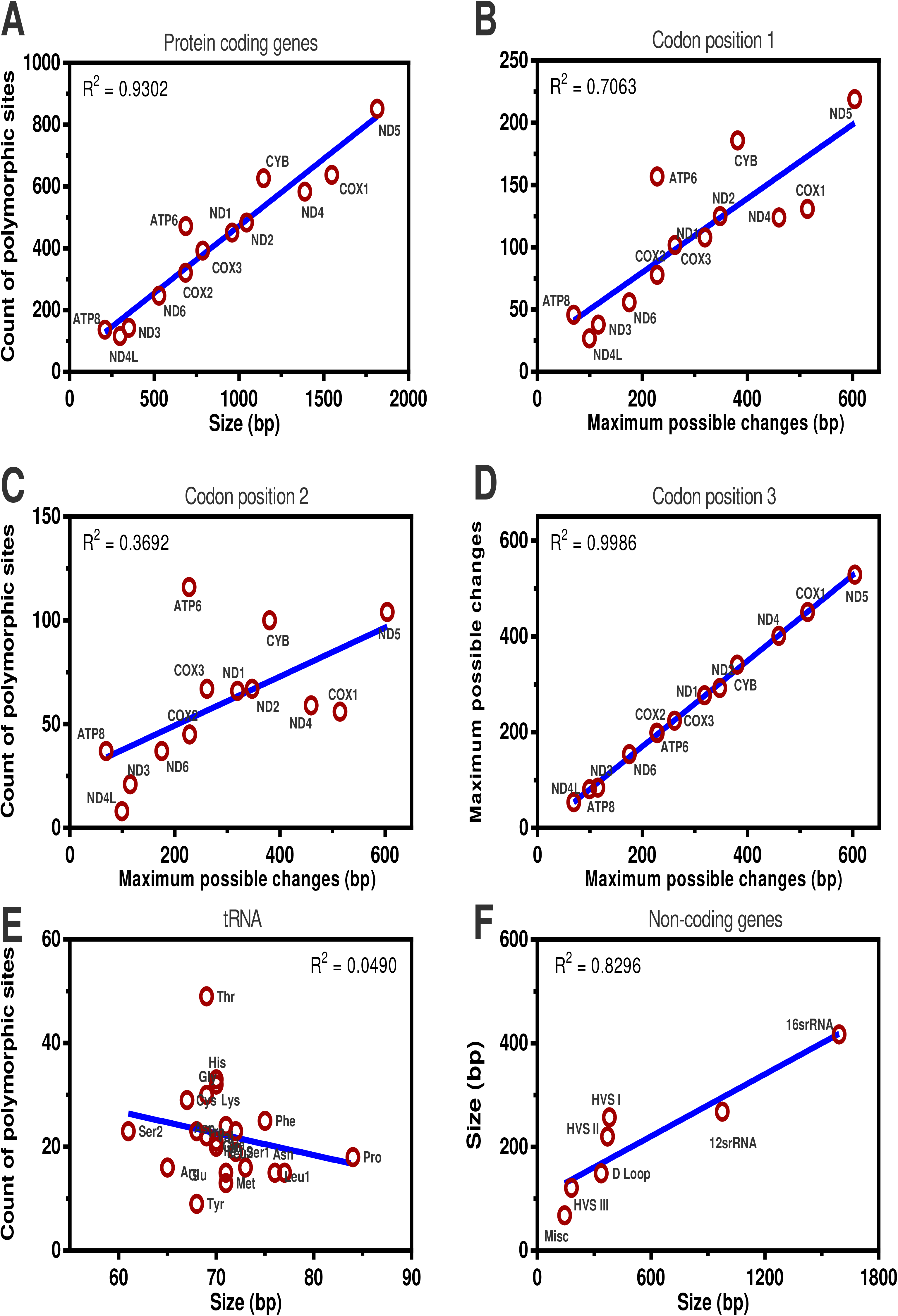
Diversity among individual genome regions. Correlation between the observed polymorphic positions and the gene size (bp) or maximum possible changes in **(A)** the 13 protein-coding genes, **(B-D)** the codon positions 1, 2 and 3 among the 13 protein-coding genes, **(E)** tRNA genes and **(F)** non-coding genes.

### 3.1. Evolution of mitochondrial co-mutations

#### 3.1.1. Co-mutations displaying intra- and inter-genomic loci biases

Analysis of pairs of co-mutations provides insight into the relationship between two distinct genome locations. Co-mutations can be formulated within a particular mitochondrial functional region (intra-loci) or between two functional regions (inter-loci). We enumerated co-mutations present among nine mt-genome functional regions. The number of polymorphic sites was normalized by the total number of co-mutating polymorphic sites in a genome group and used to construct Circos plots (Fig 4). Nine mt-genome functional regions, comprising of four oxidative phosphorylation (OXPHOS) complexes, two RNA and three non-coding regions, displayed different preferences to co-mutate with other functional regions. In particular, OXPHOS complexes I, IV and HVS functional regions have a large contribution to the overall co-mutation configuration in each network. To know more on how each functional region has contributed in forming co-mutations, we plotted the number of co-mutations in each functional region against the corresponding functional region size for intra- and inter-loci (Fig 4). It was observed that co-mutations among functional regions were evenly distributed among both intra- and inter-loci in AM and AS. However, intra-loci were more evenly distributed as compared to inter-loci. Interestingly, we reported few functional regions found to be outside the 95% confidence intervals in both intra- and inter-loci (Fig 4). For intra-loci, rRNA was an outlier in all populations, HVS in AF and OC whereas COX in AM and EU. For inter-loci, HVS was an outlier in AF, EU, and OC whereas COX in AM and AS. ATP and miscellaneous regions were outliers in AM, tRNA in AS and rRNA in OC. These statistical outlier regions should have an assertive evolutionary role in a population. To explore this further, we studied how these groups were separated from each other. We calculated Frobenius distances between each pair of five co-mutation matrices and then performed hierarchical clustering. A dendrogram clearly showed the separation of five genome groups into two main branches *i.e.* {AM, AS} and {AF, EU, and OC} (Fig 5).

**Figure 4:**
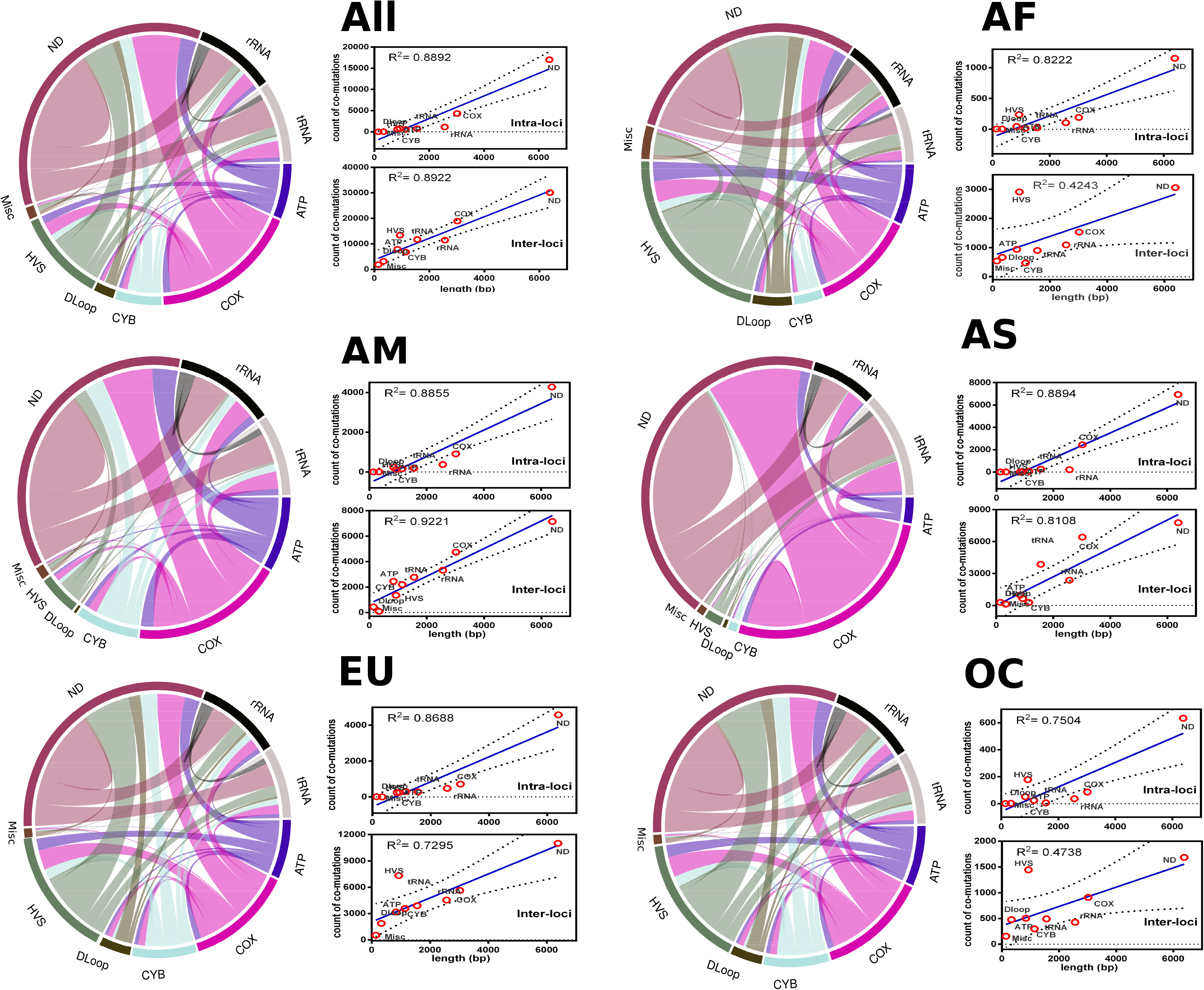
Comparison of polymorphism among genomic loci. A co-mutation configuration in the human mt-genome co-mutation network consisting of nine functional regions. These nine regions were four mitochondrial complexes (ND, COX, ATP, and CYB), three non-coding regions (DLoop, HVS, Miscellaneous) and two RNA regions (rRNA and tRNA). Links or ribbons represent the frequency of CO pairs between two genomic loci. The four functional regions make the mitochondrial oxidative phosphorylation machinery. In large part, mtDNA-specified proteins are components of respiratory complexes: Complexes I (NADH dehydrogenase), Complex III (cytochrome c), Complex IV (cytochrome c oxidase) and Complex V (ATP synthase). The regression line is shown in blue (rigid) colour whereas 95% confidence interval is shown with black (dotted) colour. Circular maps were constructed using the *rcirclize* package in R.

**Figure 5:**
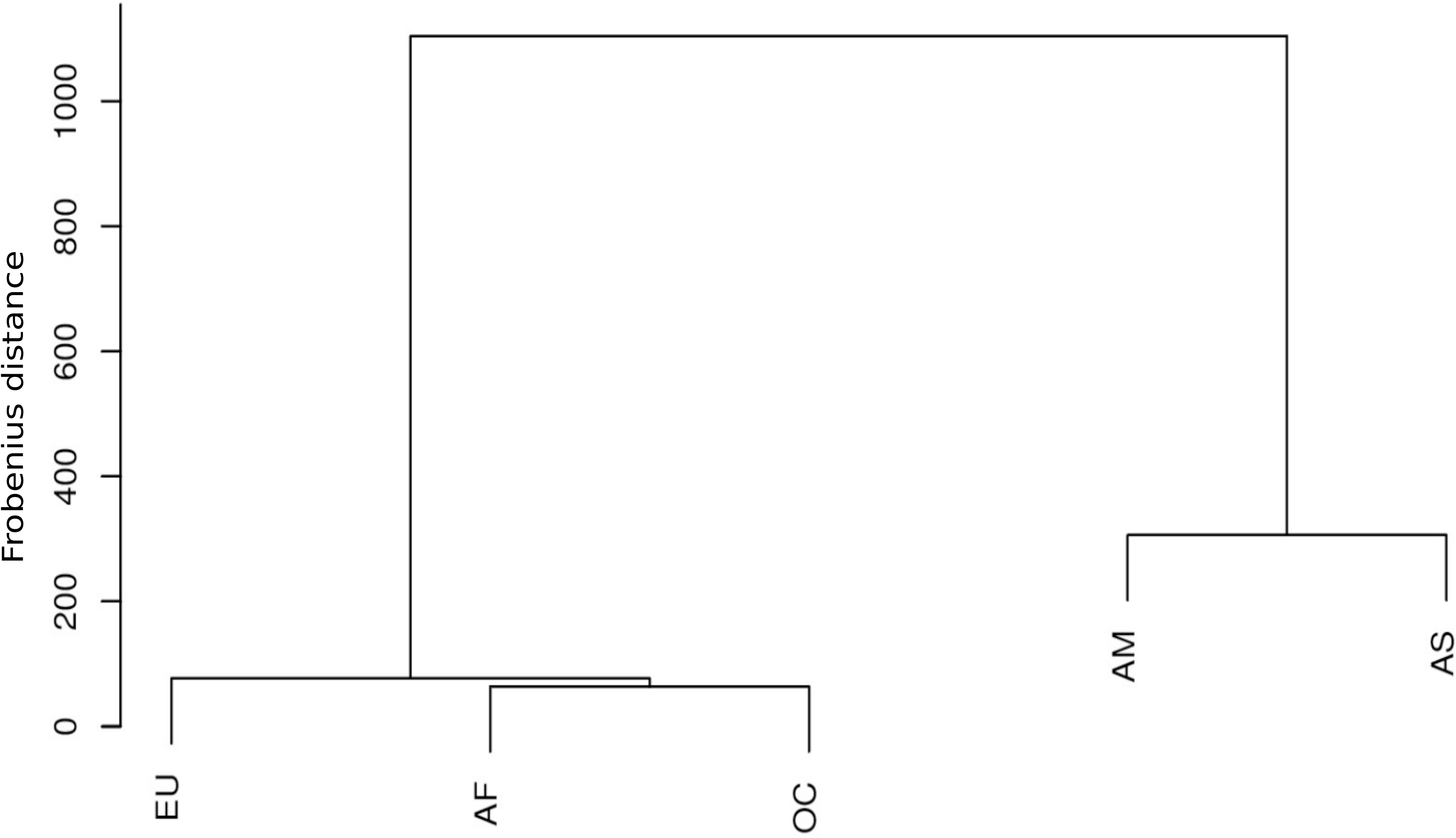
Relationships among genome groups. These relationships are predicted based on polymorphisms shown by their functional genomic loci. Five genome groups were classified as two main branches of the dendrogram *, i.e.* {AM, AS} and {AF, EU, and OC}. Here, Frobenius distance between co-mutation configuration matrices of different genome groups were used to define height of dendrogram. The branch separations shown in plot supports the routes of human migrations earlier discovered using global mt-genome mutational phylogeny. In particular, Asian haplogroup M and European haplogroup N arose from the African haplogroup L3 (Wallace, et al., 1999). Haplogroup M gave rise to the haplogroups A, B, C, D, G, and F (Wallace, et al., 1999) in which Haplogroups A, B, C, and D populated East Asia and the Americas. In Europe, haplogroup N led to the European haplogroups H, J, T, U, and V (Torroni, et al., 1996) whereas Haplogroups S, P, and Q are found in Oceania (Ruiz-Pesini, et al., 2006).

To investigate global level co-mutation preferences between functional regions, we analyzed unique co-mutations from all the genome groups. Fewer co-mutation pairs were formulated among intra-loci than inter-loci. This relationship between co-mutations and the spatial proximity is shown to be conserved in the mt-genome since all 13 protein-coding genes formed many interactions with OXPHOS complexes (Wong, et al., 1975; Thompson, et al., 1994). However, co-mutation pairs formed among OXPHOS complex I or ND genes which make 38% of total mt-genome participated in 31% of inter-loci comutations but only 13% of intra-loci co-mutations. Both D-Loop and all three hypervariable regions displayed a tendency to co-mutate with almost all other mt-genome loci (Fig 4). The rRNA genes make-up 15% of total mt-genome but they participated in only 9% of co-mutating sites. All 22 tRNA genes, which make 9% of total mt-genome, participated in 10% of co-mutating sites. Overall, co-mutations dispersed among mt-genome functional regions showed that formation of co-mutations was driven mainly by local preferences within each group. Furthermore, to investigate whether the identified co-mutations can be mapped with previously known potential disease alleles, we extracted information about potential disease alleles from Mitomap (Ruiz-Pesini, et al., 2006), OMIM (Hamosh, et al., 2005), and COSMIC (Bamford, et al., 2004) databases. There were 5, 74, 34, 8, and 10 co-mutations mapped for AF, AM, AS, EU, and OC genome groups respectively. The full list of co-mutations mapped with potential disease allele is given in File S3 and Table S6.

#### 3.1.2. Co-mutation networks exhibited similar network properties

Pair-wise co-mutations were not sufficient to fully reveal the underlying structure of functionally related nucleotide positions. As described in Fig 1, a co-mutation network was constructed for each genome group where polymorphic sites forming co-mutations constituted nodes, and edges represented co-mutating nucleotide positions. All five networks exhibited high average clustering coefficient, *〈C〉* values (Table 2), suggesting that the nodes of these networks are densely connected. Most real-world networks, particularly social networks, characterized by high *〈C〉* value, suggesting movie actors tend to create tightly knit groups by high compact ties (Sarkar, et al., 2016). In addition, all five networks displayed a highly negative degree-degree coefficient (*r*) (Table 2), suggesting that co-mutation networks were dis-assortative where high degree nodes, on average, prefer to link to low degree nodes (Newman, et al., 2003). Many biological and social networks have negative *r* values, suggesting that lack of a high degree node in a disassortative network has a large effect on the connectedness of the network (Newman, et al., 2003; Shinde, et al., 2015; Sarkar, et al., 2016). Overall, co-mutation networks have shown both the properties of high clustering and disassortative nature. This suggests the presence of dense subgraphs within the network and the presence of hierarchical structures. To explore more about the local interaction patterns in co-mutation networks, we investigated module structures within these networks.

**Table 2:** Data statistics and the properties of final co-mutation networks. Here, *α*, *N*_*P*_, *N*, *N*_*C*_, *〈K〉*, *〈〉*, *r*, *Q* represent the co-mutation frequency, number of polymorphic sites, number of nodes, number of edges, the average degree, the clustering coefficient, the assortativity coefficient and modularity coefficient for both real-world co-mutation networks and random networks, respectively. All five networks were sparse, disassortative and modular in nature. Network statistics of the largest connected component and the disconnected components are given in S1 Table and S2 Table. The number of nodes and edges forming final co-mutation networks were found to be different for each genome group. 1000 degree sequence preserved random networks are constructed for comparison of modularity in each co-mutation network and standard deviation was found to be less than 0.002 in average *Q* values of corresponding random networks.

| Network | $\alpha$ | $N_P$ | $N$ | $N_C$ | $\langle K \rangle$ | $\langle C \rangle$ | $r$ | $Q_{Real}$ | $Q_{Rand}$ |
| --- | --- | --- | --- | --- | --- | --- | --- | --- | --- |
| AF | 0.99970 | 3716 | 2310 | 13721 | 12 | 0.33 | -0.36 | 0.51 | 0.20 |
| AM | 0.99959 | 3581 | 2412 | 30283 | 25 | 0.31 | -0.61 | 0.22 | 0.10 |
| AS | 0.999854 | 5405 | 2293 | 32705 | 29 | 0.22 | -0.18 | 0.21 | 0.09 |
| EU | 0.999760 | 4557 | 2456 | 47952 | 39 | 0.18 | -0.62 | 0.30 | 0.07 |
| OC | 0.998800 | 1565 | 1208 | 7304 | 12 | 0.54 | -0.25 | 0.54 | 0.21 |

### 3.2. High cohesiveness and hierarchical organization of co-mutation communities

Is real-world network organization driven by the non-random character, at least to some extent, by modules present in the network? If this is the case, it is expected that modules would be overrepresented in original co-mutation networks compared to their counterparts such as random networks of the same size (Prill, et al., 2005). To test this, we generated random networks, referred to as a configuration model, with the same degree sequence as the original co-mutation network (Csardi, and Nepusz., 2006). Random networks lack organizing principles, therefore, the presence of modules in a random network is determined by the density of edges (Itzkovitz, et al., 2003).

The major challenge for identifying modules in a hierarchical organization is to decide the depth to decompose the network, as the Louvain algorithm fragments networks and subsequently modules until it finds the greatest partition (Meunier, et al., 2009). In order to avoid large numbers of smaller modules (size 2), the size of the second largest connected component was used to decipher submodules among each hierarchy of parent modules. The size of the second largest connected component was 11, 8, 9, 6, and 12 for AF, AM, AS, EU, OC genome groups respectively. We calculated the modularity coefficient (*Q*) for five final co-mutation networks and also for corresponding random networks (Table 2). *Q* value ranges between −1 and 1, where it takes positive values if there are more edges between same-group vertices than expected, and negative values if there are less (Blondel, et al., 2008). We tested the hypothesis that the average *Q* of random networks equals that of the co-mutation network. *Q* value was clearly reduced in the randomized networks (t-test, p *<* 0.001, for all co-mutation networks), relative to the original data, indicating that our results on real-world co-mutation networks were not trivially reproduced in random networks. A high *Q* value will manifest if networks are modular in nature. There were 557, 571, 552, 622, and 227 modules obtained for AF, AM, AS, EU, and OC genome groups respectively. The full list of modules is provided in S2 File.

In these networks, small sized modules (size less than 20) were predominant alongside one or two large sized modules *i.e.* AF (size of 119), AM (270), AS (217 and 216), AS (294) and OC (104) (S5 Fig). This signature of large-sized modules found in co-mutation networks was not displayed by corresponding random networks. Interestingly, large sized modules were only comprised of polymorphic sites from non-coding regions (except in OC). Similarly to co-mutations, we also noted that polymorphic sites among each module could be from any of mt-genome loci. For example, in the OC population, module 59 had polymorphic sites only from *COX1* gene, whereas module 3 had all polymorphic sites from different genes (S2 File). We noted that protein-coding functional regions have a predominant role in the formation of modules (S7 Table and S8 Table). Particularly, ND and COX participated in *>*65% and *>*40% of modules in each of the five networks, respectively. Additionally, we also observed a total of 391 modules out of a total of 2529 modules where all polymorphic sites in the module were from a single functional group. Such mono-functional region modules were also prevailed by ND and COX functional regions, 70% and 14% of total mono-functional region modules, respectively (S7 Table).

### 3.3. Modules of co-mutating polymorphic sites indicate ancestral relationships

To investigate if the modules identified from the analysis of the network structure were evolutionarily related, we examined polymorphic sites in the individual modules for ancestral alleles from the Reconstructed Sapiens Reference Sequence (RSRS). If a non-RSRS allele was present in more than 1% of samples in a genome group, we termed it an ancestral-variant allele. Thus, we assigned ancestral-variant information to all of the network modules and noted three distinct types of modules (Fig 6), which are explained in following

In the first and most common (more than 90% of total modules), all polymorphic sites were closely related to ancestral alleles (Table 3) and we termed them ancestral allele modules. All the polymorphic sites in these modules had ancestral alleles (or non-RSRS alleles present in *<* 1% of samples). Ancestral alleles were reported to be common throughout human mt-genome tree (Ruiz-Pesini, et al., 2006) and were also observed in large numbers in our genome group data (Table 3). In the second type of module, all the polymorphic sites were ancestral-variant alleles. We termed them as ancestral-variant modules and were of our particular interest because all polymorphic sites among these modules consist of the evolved character from RSRS. Ancestral-variant modules were observed the least out of three types of modules, both in terms of module count and the number of polymorphic sites present in these modules (Table 3). In the third type of module, polymorphic sites in a module were a mixture of ancestral and ancestral-variant alleles, and we termed them mixed modules. The polymorphic sites among these modules were hypothesized to have recently diverged. Mixed modules comprised of the large-sized modules, therefore even though the module count was found to be lower, these mixed modules still possessed a higher number of nodes (Table 3). We also confirmed that count of each module type in all five co-mutation networks is significantly different than those of corresponding random networks (t-test, p *<* 0.0001; Fig S8, S9, S10). In addition, ancestral-variant modules were difficult to produce in random networks.

**Table 3:** Statistics of modules and ancestral lineage polymorphism. Count of modules among each genome group and percentage of nodes participating in those modules (brackets) is given. Mixed modules observed to the confined of the largest size modules. 1 referes ancestral lineage polymorphic sites (ALPS)

|  | AF | AM | AS | EU | OC |
| --- | --- | --- | --- | --- | --- |
| *Modules* |  |  |  |  |  |
| Ancestral allele modules | 501 (79%) | 529 (76%) | 488 (69%) | 583 (75%) | 205 (86%) |
| Ancestral-variant modules | 26 (4%) | 12 (2%) | 11 (1%) | 12 (1%) | 12 (5%) |
| Mixed modules | 30 (17%) | 30 (22%) | 53 (30%) | 27 (24%) | 10 (9%) |
| Total modules | 557 | 571 | 552 | 622 | 227 |
| *Ancestral lineage polymorphism* |  |  |  |  |  |
| ALPS<sup>1</sup> | 163 | 192 | 136 | 177 | 122 |
| Modules with atleast one ALPS | 45 | 60 | 40 | 49 | 70 |
| Modules with >1 ALPS | 16 | 25 | 16 | 13 | 24 |
| Modules with all ALPS | 8 | 10 | 5 | 5 | 10 |

**Figure 6:**
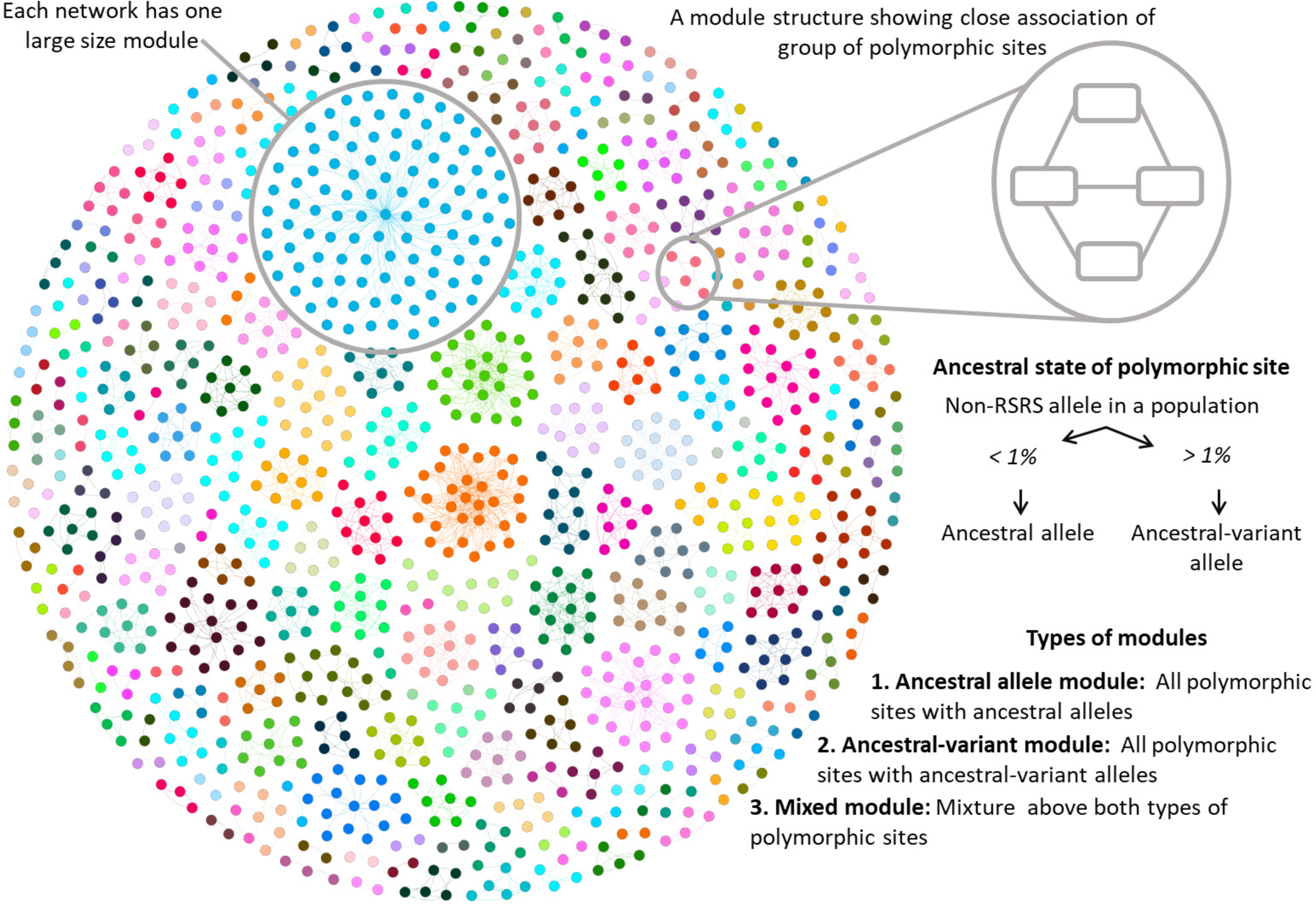
Identification and characterization of network modules. Network modules are identified using Louvain modularity algorithm. Here, the network view of modules is shown for OC network. Each network comprised of one large size module. Each polymorphic site in a module was classified as an ancestral or ancestral-variant polymorphic site. Further, modules are classified into three categories: ancestral allele module (ancestral polymorphic sites), ancestral-variant module (ancestral-variant polymorphic sites) and mixed module (both ancestral and ancestral-variant polymorphic sites).

Modules were mapped to all known haplogroups, which showed that each polymorphic site contributed to one or many haplogroups, and thus entire module structure can be related to a single mt-genome haplogroup (S2 File). Further, we investigated the relationship between modules corresponding to ancestral haplogroup lineage markers (or top-level haplogroups). Information of ancestral lineage markers was taken from the Mitomap database, and polymorphic sites among each module were mapped to ancestral lineage markers. These ancestral lineage markers were observed to participate in the formation of entire module structures, and there were a total of 38 such modules structures obtained (Table 3; File S1). Out of the observed 38 modules, where all nodes were ancestral lineage polymorphic sites, 23 were ancestral-variant modules, 13 were ancestral modules, and two were mixed modules. Since all polymorphic sites among these 38 modules were the ancestral lineage markers, it would be reasonable to say that not only sub-level haplogroups but also top-level haplogroup markers have shown a tendency to be associated to each other.

## 4. Discussion

We used comparative genome analysis to investigate 24,167 mt-genomes and devised a network model comprising pairs of co-mutating nucleotides over the human mt-genome. The method presented here provides a perspective on epistatic interactions using only sequence information as well as serves as a comparative tool to understand intra-species variations. Our study showed the presence of heterogeneity in both epistatic mutations and functional modules across investigated genome groups.

The comparison of observed polymorphisms with gene size clearly showed two essential features in providing maximum functional level diversity with the minimum level of genomic changes. First, genetic conservation at CP 2 but not at CP 3, was key to providing a protein diversity of mt-genome complexes. Second, the restriction of mutations in tRNA and rRNA. These two observations of biases against mutations at CP 2 and RNA genes were earlier reported by Pereira et al. with 5140 human mt-genome sequences (Pereira, et al., 2009). The similar biases of CP 2 and tRNA genes were also reported among mt-genomes of other primates including Macaca, Papio, Hylobates, Pongo, Gorilla, and Pan whereby the strength of selection was determined in each lineage by the ancestral state of each codon position (Kivisild, et al., 2006). Among non-coding genes, all three HVS regions have displayed a higher level of polymorphisms, whereas genes of rRNA and tRNA have shown lower levels of polymorphism. Our study, apart from providing the detailed enlisting of diversity present among five genome groups, reiterated that both codon level mutation bias and restriction of mutations among RNA genes were more evident at the subpopulation level despite infrequently reported to be at a global level. Furthermore, given the ubiquitous variation in mt-genome, genetic flexibility may have evolved as a mechanism to maintain OXPHOS under a range of environments.

As well as biases against polymorphisms at CP 2 in protein-coding genes, our analysis indicates other biases with co-mutations. First, our results with intra- and inter-loci preferences clearly suggested the dominance of polygenic mutations in the human mt-genome. These polygenic mutations are the outcome of a highly constrained organization of OXPHOS complexes (Fonseca, et al., 2008) and also due to protein-protein interactions of the mtinteractome (Schweppe, et al., 2017). Second, our analysis highlights regions of the mt-genome which are rich in co-mutations and thus suggests the presence of epistasis. In particular, a large portion of COX genes co-mutate in AS and AM populations whereas in AF, EU and OC populations, there was greater co-mutation bias in functional regions of HVS. Although mitochondrial genome epistasis is largely described in the context of mitochondrial-nuclear interaction due to the closed assembly of OXPHOS complexes (Fonseca, et al., 2008; Picard, et al., 2018; Connallon, et al., 2018; Schweppe, et al., 2017), there are many reports describing the presence of mitochondrial-mitochondrial epistasis, for example, shared family features in Han Chinese family (Wang, et al., 2015) and homoplasy guided by mt-tRNA genes (Moreno-Loshuertos, et al., 2011). Furthermore, few co-mutations were found to be mapped with previously known potential disease alleles, implicating association of disease phenotypes within mt-genome sequences. Mitochondrial epistasis has been reported for its role in mitochondrial diseases (Morrow, and Camus., 2017; Smith, and Lusis., 2002; Pritchard, et al., 2010; Schrider, and Kern., 2017). We also note that these individual mapped co-mutations do not possess potential disease alleles in other genome groups. In view that mt-genome lineages are functionally different, it reflects that the same epistatic interaction can be advantageous in one environment and might be maladaptive in a different environment (Mishmar, et al., 2003). Hence, lineage-defining haplotypes could be contributing to bioenergetic disorders as the migration takes place (Mishmar, et al., 2003). Third, similar to polymorphic sites, co-mutations also showed biases at the subpopulation level. Genome group-wise comparison of co-mutations associated with mt-genome functional regions has helped in classifying these five human subpopulations into two prominent groups *i.e.* {AF, EU, OC} and {AS, AM}. This result was supported by a global mt-genome mutational phylogeny (Ruiz-Pesini, et al., 2006) showing the routes of human migrations (Fig 5). Overall, variations probed by epistatic interactions have provided local preferences among different mt-genome loci. These local preferences might have helped in not only forming the closed-assembly of OXPHOS complexes but also classifying subpopulations.

In our network model, the emergence of sparse networks was not a smooth, gradual process: the very dense largest connected component collapsed into a sparse largest connected component through a sudden change in the *α* curve (Fig 2). For all five genome groups, we encountered such a distinct phenomenon. A similar critical phenomenon was first observed by Erdös and Rényi through their random network model where the isolated nodes and tiny components observed for small *〈K〉* would collapse into one largest connected component (Erdos and Rnyi., 1960). Interestingly, the nature of discontinuous transformations was earlier reported in biological networks (Fontana, and Schuster., 1998; Liu, et al., 2012) and have been hypothesized that biological processes follow discontinuous transformation during their evolution (Fontana, and Schuster., 1998).

We selected edges for inclusion in co-mutation networks based on their best fit to a network sparseness. Sparseness is one of the essential properties of biological networks since links are more difficult to create due to the evolutionary cost involved in forming more links. It is well known that co-mutational events are very selective and require a group of cooperative supporting mechanisms (Du, et al., 2008). Previously, modules of highly correlated genes were identified using similar edge-filtering based methods like weighted correlation network analysis (Jackson, et al., 2018). Network sparseness or similar data-driven approach avoids arbitrary selections of network edges and provides a uniform rationale that can be implemented to generate co-mutation network structures across different genome datasets. Therefore, it was reasonable to choose an *α* value where a network should have both the lowest value of *〈K〉* and the largest component with a higher count of nodes.

There was clear evidence for hierarchical modularity in our genome datasets, and the modular structure of the networks at all levels of the hierarchical patterns was reasonably similar across genome groups, suggesting that mt-genome functional modularity is likely to be a replicable phenomenon. A combination of co-mutations at different mitochondrial regions, that are closely linked, tend to be inherited together. This study provides a complete listing of the current knowledge of mt-genome variation in the human population, also with respect to their higher level associations with hierarchical modules. Every set of co-mutations found to originate from and remain part of a preceding single group of co-mutations. This nested hierarchy suggests the conservation of ancestral as well as inherited co-mutations throughout human lineages. Similar hierarchical modularity in brain network was related to functional regions in the brain and sub-set of brain functions have been reported to be associated among each hierarchy (Meunier, et al., 2009). Modularity is one of the main features of co-mutation networks, and evolutionary processes may favor the emergence of modularity by a combination of structural and functional preferences in forming molecular interactions (Clune, et al., 2013). Therefore, it was reasonable to say that evolutionary processes may favor modularity by allowing both the specificity and autonomy of functionally distinct subsets of genomic positions. Overall, we demonstrated that molecular changes, such as mutations, were not randomly distributed across the genome, but instead concentrated within modules. In this sense, the concentration of genomic positions within modules provided a way to understand module integration, favoring distinct functional roles developed by genomic positions in distinct modules. In the human mt-genome, modules were associated with mitochondrial subcomplexes that act in distinct steps of the electron transport assembly and function. Thus, the closed assembly of mitochondrial complexes might favor the emergence of highly integrated genomic subunits, in which effects of pairwise interactions may also activate indirect effects on non-interacting genomic positions associated with the same function (Fonseca, et al., 2008). In addition, the OXPHOS system is intrinsically incapable of evolving to a fixed and general optimum state; therefore, both functional and genetic heterogeneity plays a vital role in providing robustness to the evolving OXPHOS system (Enriquez., 2016). Based on these results, we would expect that genome positions connecting modules were more conserved across evolution or, at least, less prone to failures that alter their function.

It was expected that module level associations would reflect evolutionary relationships between underlying genomic positions as each module consisted of ancestrally similar genomic polymorphisms. Our results added that the distinction between ancestral and ancestral-variant mitochondrial polymorphisms was clear when the entire module was made up of ancestral-variant polymorphic sites. However, a large number of modules (more than 90% of total modules) were made-up of ancestral polymorphic sites. In addition, the large number of nodes in mixed modules were of ancestral origin (S2 File). Using the list modules of all five networks, it would be reasonable to assert that contemporary mt-genome nucleotide bases most closely resembled the ancestral state and very few of them were ancestral-variants. This observation was in agreement with previous studies which found co-mutation among nucleotide positions to be higher between genetically similar taxa (Chaffron, et al., 2010). This fact was widely observed in our data as both sub-level, and top-level haplotype markers were associated with each other in a closed group of network modules. Furthermore, mitochondrial polymorphic positions adapt from ancestral state to ancestral-variant state (Keightley, and Jackson., 2018), is also demonstrated by a larger count of transient state mixed modules. Overall, these evolutionarily closed associations suggest that interactions between nucleotide positions might evolve within genetically related genomic polymorphic positions (more likely of having similar functionality) responding to intra-species biases (Du, et al., 2008).

Understanding the formation of the network would require an extension of the described approaches. Here, we used simplistic information possessed by each genome position in terms of their underlying ancestral markers. Previously, this ancestral marker information has been used in order to define taxa (precisely haplogroups) in mitochondrial phylotrees which have provided the exact mapping of mitochondrial signatures to infer the routes of human intra-species diversification events (Nakatsuk, et al., 2017; Derenko, et al., 2001). Similarly, our co-mutation modules have provided a detailed listing of mitochondrial co-mutations which were ancestrally associated together.

Consistent with the proposed importance of mt-genome variation in human adaptation (Wallace., 2015), regional haplotypes are generally founded by one or more functionally significant polypeptide, tRNA, rRNA, and control region variants. These variant traits, beyond being retained in the descendant population, manifest functional signatures about affecting phenotypic variations in the subpopulation. Particularly, a list of singular events involving both mt-genome variation and epistatic interactions have been evaluated in terms of affecting phenotypic variation in metabolism, fitness, and life-history traits (Wallace., 2015; Shlush, et al., 2008; Li, et al., 2015). Hence, it is intuitive to observe epistatic patterns of both genotype and phenotype variants along with life-history traits at the subpopulation level. However, at the between-population level, the evidence in support of the relationship between human mt-genome variation and the metabolic rate is compelling.

## 5. Conclusion

We constructed and investigated human mt-genome co-mutation networks of continents using a combined framework of genomics and network theory. Our principal result was that mitochondria undergo substantial levels of co-mutational biases. Codon-level mutation bias, particularly at CP 2, and restriction of mutations in RNA genes was also evident at the continental level, which was earlier only reported in the global human population. The analysis highlighted regions of mt-genome rich for co-mutations and thus suggested the presence of epistasis. In particular, a large portion of COX and ND genes found to be co-mutated in AS and AM populations whereas in AF, EU, and OC populations, there was greater co-mutation bias between regions of HVS and ND, thus our networks identified differences in co-mutation bias between human populations. It was of great interest to investigate and verify different co-mutation patterns of various geographical regions. Importantly, we deduced hierarchical modular structures formed within co-mutation networks. Downstream analysis of these modules suggested that contemporary human population are dominated by ancestral states. In addition, ancestral-variant module structures are found to be in a lesser number, and such modules have found to be difficult to produce in corresponding random networks. The analysis presented here can be extended to study the complexity of mt-genome evolution by forming various geographical groups as well as to understand alterations in personal traits leading to complexity in mt-genome evolution.

## Acknowledgments

We are grateful to our the editor and reviewers for helping us to improve the manuscript. PS acknowledges Inspire fellowship [IF150200] from the Department of Science and Technology (DST), Government of India. SJ thanks the support by grant of DST [EMR/2016/001921], Government of India. RKV acknowledges CSIR-NET fellowship [Roll No.: 305089] from CSIR, Government of India.

## Data Availability Statement

All data sources and related information is given in the manuscript and associated supplementary material files.

## Supplementary materials

**S1 File. Information of genome samples considered in the study.**

**S2 File. Information of modules identified in the study.**

**S3 File. Information of co-mutations consist of potential disease alleles (PDA).**

**Table S1:** Network properties of Largest connected component (Lcc).

**Table S2:** Network properties of all disconnected components together except Lcc.

**Table S3:** Level-wise community detection in five co-mutation networks.

**Table S4:** Comparison of polymorphism *α*_*pre*_ and *α*_*post*_.

**Table S5:** List of hub nodes among co-mutation networks.

**Table S6:** List of co-mutations consist of potential disease alleles (PDA).

**Table S7:** Distribution of modules having at least one polymorphic site among mt-genome functional groups.

**Table S8:** Modules comprising all nodes as ancestral lineage polymorphic sites.

**Table S9:** Proportion of modules among co-mutation networks and corresponding random networks.

**Table S10:** Proportion of nodes in modules among co-mutation networks and corresponding random networks.

**Figure S1:** Distribution of variable sites across genome groups and the distance between each genome from the reference sequence (RSRS).

**Figure S2:** Degree distribution of five co-mutation networks.

**Figure S3:** Gene-wise comparison of polymorphism before and after *α*.

**Figure S4:** Transition and transversion.

**Figure S5:** Schematic description of assortativity.

**Figure S6:** Distribution of module sizes.

**Figure S7:** Identification and characterization of network modules.

**Figure S8:** Distribution of modules count in random networks.

**Figure S9:** Distribution of modularity coefficient among random networks.

**Figure S10:** Count of different type of modules among random networks

